# Biodiversity monitoring in agricultural landscapes: Why it matters

**DOI:** 10.64898/2026.02.28.708698

**Authors:** Luciano M. Verdade, Rafael A. Moral

**Affiliations:** Wildlife Management Consultancy, Campinas, SP, Brazil; Department of Mathematics and Statistics, Maynooth University, Maynooth, Ireland

**Keywords:** additionality paradox, carbon and biodiversity credits, guardianship, multifunctional landscapes, standardized monitoring protocol

## Abstract

Although agricultural expansion and intensification have caused extensive biodiversity loss, agricultural landscapes remain central to global conservation outcomes. No country can conserve its entire biota exclusively within conservation units, even under ideal management. Consequently, biodiversity conservation, ecosystem service provision, and food security increasingly depend on how agricultural landscapes are managed. We argue that quantitative, standardized biodiversity monitoring is paramount for aligning agricultural production with biodiversity conservation. We highlight a structural limitation in current crediting frameworks that renders long-term guardianship economically invisible relative to post-disturbance recovery. Using empirical evidence from regenerated forests, we show that guardianship can deliver substantially greater carbon benefits than additionality alone. Together, these perspectives provide a framework for integrating biodiversity conservation and agricultural sustainability in multifunctional landscapes.

---

Agriculture is still the basis of what we call civilization with a deep influence in human history, culture, socioeconomics, and even religion (*1*). However, historical land use changes caused by agriculture brought a plethora of environmental impacts which resulted in population decline and local extinction of many wild species due to a combination of habitat loss, environmental contamination, and the introduction of exotic competitors, parasites, and pathogens. However, despite all its negative impacts, agricultural landscapes can be relevant to the conservation of a significant part of biodiversity as no country is able to provide conservation for its whole biota only in conservation units (e.g., national parks and ecological reserves) (*2*). In a land-sharing approach (*3*), multifunctional landscapes can be considered as those which keep the main mission of biological production with a secondary – but fundamental – mission of biological conservation.

Most of the wild species of animals and plants that make use of agricultural landscapes are generalists (*4*). However, useful, damaging, and even endangered species can also be found in these systems (*5*). A pragmatic yet effective approach suggests that there are only four alternatives to human intervention in the field of wildlife management: increase a depleted population, decrease an excessive population, use a valuable population on a sustainable basis, or ‘do nothing but keep an eye on it’ (*6*). These alternatives correspond to the four applied evolutionary ecology-based management actions, respectively: biological conservation, damage control, sustainable use, and monitoring. Monitoring is not surprisingly the major demand both in pristine and anthropic landscapes as most species are not endangered, damaging, or useful, but can become one of those because of novel and dynamic anthropic pressures (*7*). Notwithstanding, biodiversity monitoring in agricultural landscapes still has four main conceptual, technical, and societal limitations.

First, the attribution of “non-habitat” status to anthropic environments should be tested as a hypothesis rather than be taken as an assumption (*8*). By doing the latter, species’ adaptive use of managed landscapes is overlooked and ecological functions taking place outside native areas are masked. Instead, if the “non-habitat” status of human-altered landscapes is treated as a testable hypothesis, this would allow policy frameworks to incorporate evidence of species persistence and resource use, as well as ecosystem functions. Second, indicator species tend to be chosen based on a specialist-specific approach, whereas that should be done based on a more effective problem-oriented approach. The indicators should be chosen to detect specific pressures, policy targets, or ecosystem services, thereby improving their interpretability and relevance in monitoring programs. This, in turn, would enable a more effective assessment of anthropic interventions. Third, sampling methodologies developed for temperate pristine environments are still widely used in tropical anthropic regions. These designs may fail to capture the intricacies of tropical systems, therefore yielding a misrepresentation of impacts and trends. This weakens the evidence base for policy in areas of rapid land-use change, especially given the current expansion of agriculture in these regions. Finally, in regions where land-use policy is based on a land-sparing approach, governance tends to be ineffective to promote biodiversity conservation initiatives in agricultural landscapes (*3*). Land-sparing policies tend to concentrate conservation efforts within the protected areas, while the biodiversity outcomes in agricultural landscapes are weakly regulated (or even neglected). This governance gap, coupled with limited enforcement capacity, further reduces the effectiveness of conservation efforts by ignoring connectivity and cumulative impacts at the landscape scale.

## A standardized protocol for multifunctional landscapes

Agricultural fields cover approximately 32.6% (47.81 Mkm^2^) of the land on Earth, whereas protected areas cover approximately 16.9% (24.75 Mkm^2^) (*9, 10*), with a varying proportion across continents (Fig. 1a). However, in areas with a land-sharing approach, agricultural landscapes maintain relevant fragments of native vegetation (*11*). Brazil provides a particularly illustrative case (Fig. 1b), with protected areas encompassing 31.2% of the territory (2.64 Mkm^2^) and agricultural lands occupying a comparable 28.0% (2.37 Mkm^2^). The Brazilian Forest Code demands at least 20% of native vegetation maintained in rural properties based on their original as Legal Reserve (LR). In addition, Areas of Permanent Protection (APP) are required to partially occupy riparian areas and ravines (*12*). If effectively enforced, APP and LR would represent a significant increase of native (re)vegetation in rural properties. This would improve biological conservation and ecosystem service provision without necessarily compromising biological production, which could be the basis for establishing multifunctional landscapes in Brazil and other countries.

**Fig. 1.**
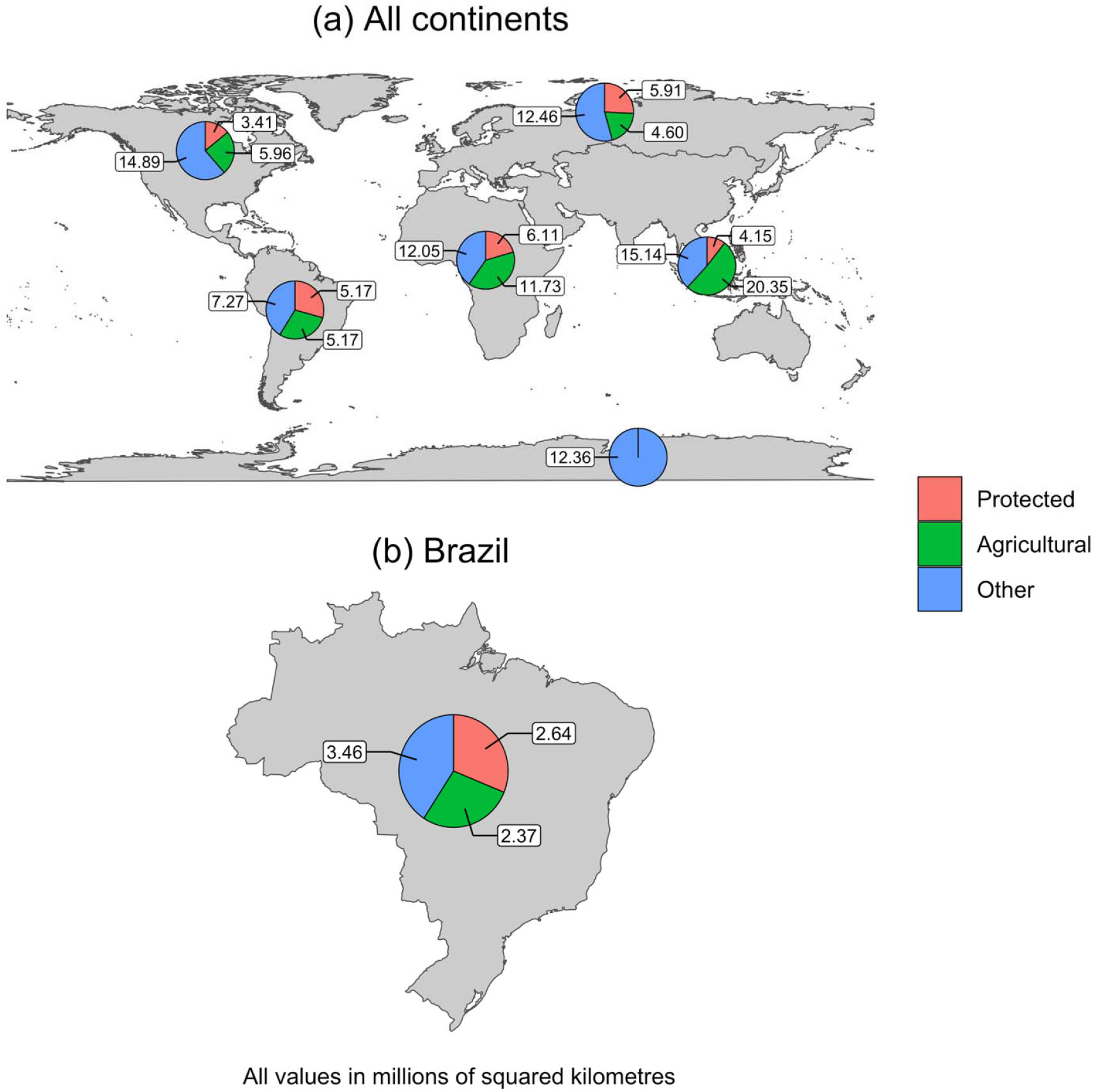
Land use: agricultural vs. protected areas. **a**, Protected, agricultural and other land use areas for the North America, South America, Antarctica, Europe, Africa and Asia-Pacific regions. **b**, Protected, agricultural and other land use areas in Brazil. All values in Mkm^2^. Sources: (*8, 9*).

For multifunctional landscapes to be effective, however, a standardized protocol for biodiversity monitoring should be established to support the decision-making process concerning both agricultural and wildlife management (*13, 2*). Such protocol should have the following characteristics:

a. Temporal scale: long-term, ideally permanent or at least for as long as agricultural activities remain;
b. Spatial scale: cross-scale, from local to regional, including a network of long-term field sites which are representative of each large agricultural sector;
c. Remote sensing: an intensive use of remote sensing should connect local-scale data derived from long-term measurements from specific field locations to the wider regional scale covered by the large agricultural sectors;
d. Spatial analyses: besides area, height of native vegetation should be included as it is correlated with volume and, consequently, with above-ground biomass of native vegetation and carbon, as well as with patterns of biodiversity and biocomplexity (*2*). This would represent a change from 2-dimensional to 3-dimensional spatial analyses;
e. Cost-effectiveness: cost-benefit relationships should be considered in research and development programs related to biodiversity monitoring technology, as cost usually limits their application on a commercial scale;
f. Indicators: metrics should include diversity of patterns and complexity of processes, focusing on the balance between natality/immigration rate and mortality/emigration rate of the species of interest;
g. Socioenvironmental certification: multifunctional landscapes should be certified to become eligible to financial carbon (C) and biodiversity (BD) credits;
h. C and BD credits should include not only additionality (from e.g., revegetation and rewilding), but also guardianship (from e.g., the conservation of areas of native vegetation in climax stage, either pristine or recovered).

## The “additionality paradox”

When implementing C and BD credit schemes, if guardianship is unrewarded, landscapes that have practically no additionality have little economic visibility, despite delivering high and sustained C and BD benefits. As a practical example, a simple analysis of mean C additionality over time across regenerated forests globally (Fig. 2a) and in the Neotropics (Fig. 2b) reveals a growth that saturates after approximately 50 years. A simple calculation of the accumulated C during the growth phase (years 0-50), where the economic value of additionality would be explicit, versus accumulated C after saturation (years 50-100), corresponding to guardianship, reveals that the guardianship period would account for 2.69 to 2.78 times more estimated C accumulation when compared to the additionality period (Fig. 2a and 2b, respectively). The lack of economic visibility of guardianship within current crediting frameworks may create an inadvertent incentive in which ecosystem disturbance followed by recovery becomes creditable, whereas long-term protection does not, as only post-disturbance gains qualify as additional. This does not imply intentional ecosystem disturbance but reflects a structural feature of current crediting frameworks that recognize recovery while rendering protection economically invisible. We refer to this structural inconsistency as the “additionality paradox”. These principles also apply to BD, insofar as both C and BD are associated with the above-ground biomass of native vegetation (*2*). Incorporating cost-opportunity approaches into C and BD markets could be a conceptually simple yet effective way to render guardianship tangible.

**Fig. 2.**
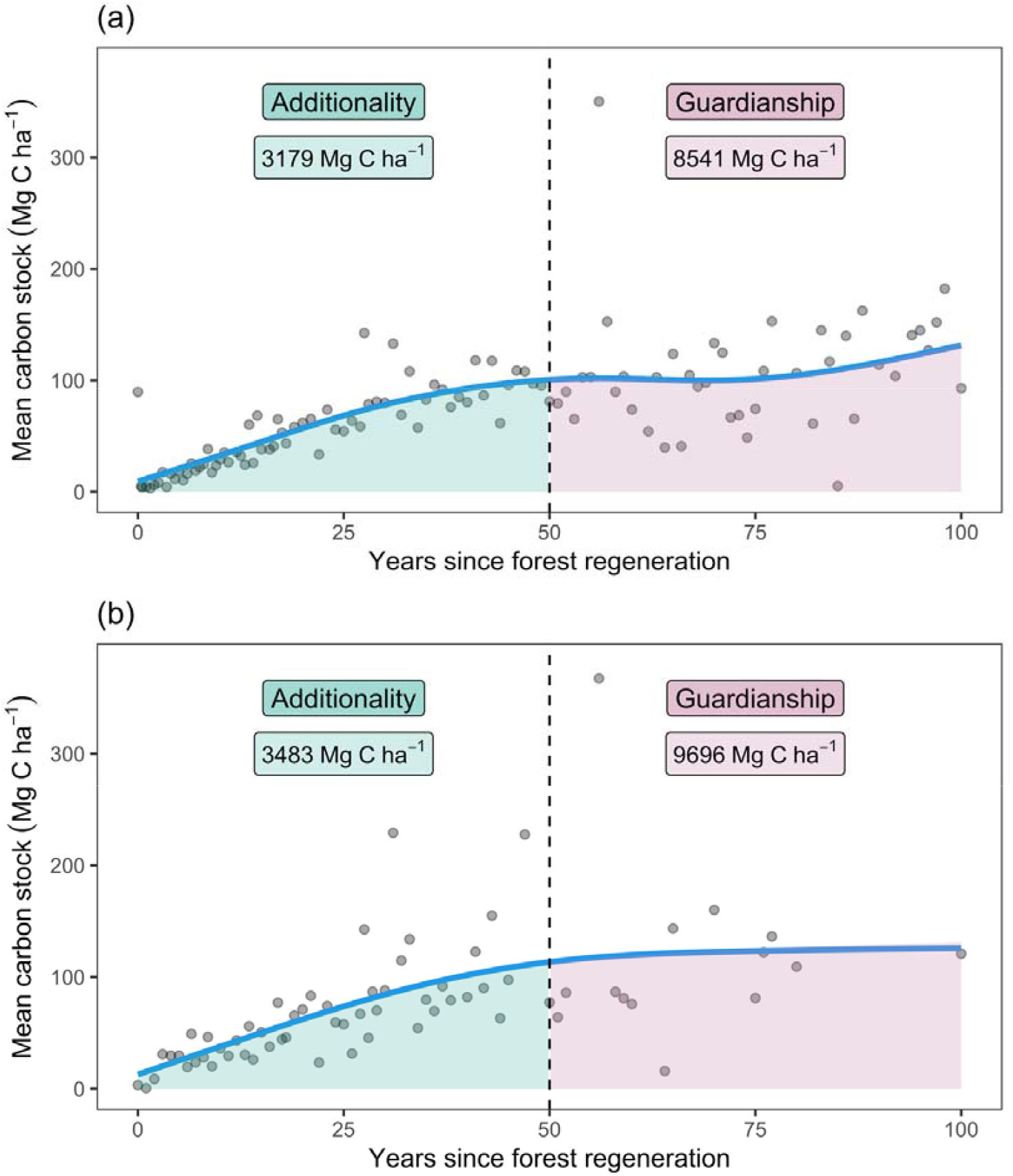
Carbon stock additionality through natural forest regeneration. **a**, Mean carbon stock across forest biomes globally, including forests in North and South America, Europe, Africa, and the Asia-Pacific regions (447 sites) versus total number of years since forest regeneration. Aboveground biomass (AGB, Mg ha^-1^) data was obtained from (*14*) and transformed to carbon stock (C, Mg ha^-1^) via the linear transformation C = 0.47 * AGB. **b**, Mean carbon stock across Neotropical forest biomes (41 sites) versus total number of years since forest regeneration. Aboveground biomass (AGB, Mg ha^-1^) data was obtained from (*15*) and transformed to carbon stock (C, Mg ha^-1^) via the linear transformation C = 0.47 * AGB. The blue curves represent predicted mean carbon stock obtained via a generalized additive model including thin-plate regression splines over time. The shaded areas represent the estimated accumulated mean carbon stock for the additionality (years 0 to 50) and guardianship (years 50 to 100) periods.

The implications of this additionality paradox extend beyond forest carbon dynamics to managed and agricultural landscapes, where long-term guardianship of biodiversity underpins ecosystem resilience and food security. The wild lineages of domesticated species of animals and plants are still fundamental to the genetic improvement of their domestic relatives against climate change and other anthropic negative impacts. Therefore, the conservation of wilderness is fundamental to provide agricultural sustainability. The recommendations above might help convert the intrinsic conflict between biological production and biological conservation into an explicit interdependence. They might also make the conservation value of agricultural landscapes tangible by the quantification of their diversity of biological patterns and complexity of evolutionary-ecological processes. And this is why monitoring biodiversity in agricultural landscapes matters.

## Author contributions

Conceptualization: LMV, RAM

Methodology: LMV, RAM

Visualization: RAM

Funding acquisition: LMV, RAM

Writing – original draft: LMV

Writing – review & editing: LMV, RAM

## Data, code, and materials availability

Open access data to reproduce Fig. 1 are available at https://archive.ourworldindata.org/20250624-125417/grapher/agricultural-land.html (*9*) and https://www.protectedplanet.net/en/thematic-areas/wdpa?tab=WDPA (*10*). Open access data to reproduce the analysis in Fig. 2 are available at https://zenodo.org/records/3983644

(*14*) (Fig. 2a) and https://datadryad.org/dataset/doi:10.5061/dryad.82vr4 (*15*) (Fig. 2b). All code is available at https://github.com/rafamoral/why_monitoring_matters.

